# Substance P receptor signaling contributes to host maladaptive responses during enteric bacterial infection

**DOI:** 10.1101/2024.06.24.599421

**Authors:** Michael Cremin, Valerie T. Ramirez, Kristina Sanchez, Emmy Tay, Kaitlin Murray, Ingrid Brust-Mascher, Colin Reardon

## Abstract

Immune responses in the intestine are intricately balanced to prevent pathogen entry without inducing immunopathology. The nervous system is well-established to interface with the immune system to fine-tune immunity in various organ systems including the gastrointestinal tract. Specialized sensory neurons can detect bacteria, bacterial products, and the resulting inflammation, to coordinate the immune response in the gastrointestinal tract. These sensory neurons release peptide neurotransmitters such as Substance P (SP), to induce both neuronal signaling and localized responses in non-neuronal cells. With this in mind, we assessed the immunoregulatory roles of SP receptor signaling during enteric bacterial infection with the non-invasive pathogen *Citrobacter rodentium*. Pharmacological antagonism of the SP receptor significantly reduced bacterial burden and prevented colonic crypt hyperplasia. Mice with SP receptor signaling blockade had significantly reduced inflammation and recruitment of T-cells in the colon. Reduced colonic T-cell recruitment is due to reduced expression of adhesion molecules on colonic endothelial cells in SP receptor antagonist-treated mice. Using SP receptor T-cell conditional knockout mice, we further confirmed SP receptor signaling enhanced select aspects of T-cell responses. Our data demonstrates that SP receptor signaling can significantly reduce inflammation and prevent host-maladaptive responses without impinging upon host protection.

## Introduction

Control of immune responses in the gastrointestinal tract is a critical determinant of health. This mucosal surface is exposed to a variety of antigens derived from commensal microorganisms; however, it can also be an entry point for pathogens. Perhaps unsurprisingly, a variety of overlapping regulatory mechanisms have evolved that allow for robust host protection, while limiting inappropriate responses to otherwise innocuous substances. Communication between the nervous and immune systems is now well established to regulate immune cell function and consequently immunological outcomes in several organ systems including the intestinal tract^1–4^. These neuroimmune interactions are aided by the colon’s dense innervation, with neurons that reside within (intrinsic) and neurons with cell bodies that are outside (extrinsic) but project into discrete portions of the intestine. Extrinsic innervation encompasses highly specialized sensory neurons that express the polymodal nociceptive receptor TRPV1, and as such can be activated by noxious stimuli such as extreme heat, danger-associated molecular patterns, bacterial components, and immune mediators produced as a consequence of inflammation^1,5,6^.

Far from simple detection of inflammation, sensory neurons are required to coordinate host protection during enteric bacterial infection. Activation of these neurons not only can induce classical reflex arc where ascending signals are coordinated in the dorsal root ganglia to drive efferent neuronal activity but can cause localized release of neurotransmitters. During *Salmonella typhimurium* infection of the small intestine, release of the neuropeptide Calcitonin gene-related peptide (CGRP) was found to exert a host protective role^7^. Similarly, we have previously demonstrated that mice with targeted depletion of peripheral sensory afferent neurons^8^, or TRPV1 knockout (KO) mice^9^, experienced increased bacterial burden and enhanced colonic pathology during *Citrobacter rodentium* infection. Although our studies suggested that CGRP did not exert a host protective role, the neuronally derived signal that coordinated host responses during enteric bacterial infection with this pathogen was not elucidated. Upon activation, nociceptive neurons release several immunologically relevant neurotransmitters in addition to CGRP, such as Substance P (SP). This neurotransmitter has long been described to not only be the mediator encoding pain but has immune effects as well^1,2,10,11^. Localized release of SP is the basis for neurogenic inflammation, where this peptide acts as a chemoattractant that can activate blood endothelial cells (BEC), causing increased expression of adhesion molecules and vascular permeability^12,13^. It is critical to note that these physiological effects are due to activation of the SP receptor where three ligands have been identified to induce receptor activation. These include not only SP and Neurokinin A, encoded by the pre-protachykinin A/Tac1 gene, but also hemokinin-1, encoded by the Tac4 gene^14–17^. In the intestinal tract, SP signaling through the SP receptor (TACR1) has a proinflammatory effect. During acute induced models of colitis, deficiency in SP, or prior ablation of sensory afferent neurons, reduced disease severity^18^. Similarly, the highly selective SP receptor antagonist CP96345 significantly reduced DSS-induced inflammation in rats^19^. Thus, SP receptor signaling can increase host-maladaptive immune responses *in vivo* resulting in immunopathology. Highlighting the complexity of neuroimmune responses and in contrast to SP receptor signaling driving host-maladaptive responses in colitis, this receptor appears to be critical to the host response during bacterial or parasitic infection. Infection with the invasive enteric bacterial pathogen *S. typhimurium* was significantly worsened with SP receptor antagonists due to reduced Th1 immune responses and mortality. This dichotomous role of SP receptor signaling suggests it has a unique role in the control of mucosal immunity, where the outcome for the host is dependent on the specific challenge or cause of inflammation.

The host immune response to *C. rodentium* is comprised of an early innate immune response, followed by the generation of CD4+ T-cells and the eventual production of antibodies^20,21^. Although required for control and eventual clearance of the pathogen, these immune responses cause significant immunopathology^22,23^. In addition to the recruitment of various immune cell types, cytokines such as IFNγ induce crypt hyperplasia which is the rapid proliferation of intestinal epithelial cells (IEC), resulting in the loss of mature secretory and absorptive cell types in the colon^22,24^.

Here, we assessed the role of SP receptor signaling on the course of disease and the immune response elicited during infection with the non-invasive enteric bacterial pathogen *C. rodentium*. Using potent and selective TACR1 antagonists, we demonstrate significantly reduced *C. rodentium* burden, and infection-induced colonic pathology. Surprisingly, antagonism of the SP receptor significantly reduced selected aspects of the host response to this enteric bacterial pathogen, with decreased colonic T-cell recruitment, and expression of critical cytokines 10 days post-infection (dpi). Analysis of the colon in these antagonist-treated and infected mice revealed significantly decreased expression of the adhesion molecule MAdCAM-1, which serves to recruit mucosal homing T-cells to the intestine. In addition to reducing the recruitment of colonic T-cells, significantly reduced IFNγ production and IFNγ+ CD4+ T-cells were observed in the colon 10 dpi in TACR1 antagonist-treated mice compared to vehicle. These effects on T-cells were further determined to not be due to changes in conventional dendritic cell (cDC) numbers or a reduced ability of these cells to present antigen, and T-cells to respond *in vivo* to an orally administered antigen. With the recent demonstration that the SP receptor is required for T-cell receptor-induced Ca^2+^ signaling^25^, we investigated the effect of T-cell-specific TACR1 deficiency during *C. rodentium* infection. Using a unique TACR1 T-cell conditional KO mouse, we demonstrated that loss of T-cell intrinsic SP receptor signaling does not reduce colonic T-cell homing, but significantly attenuates T-cell IFNγ production. Together, our data suggests that SP receptor signaling contributes to host defense through a variety of discrete cell types including T-cells. Moreover, our data demonstrate that this receptor could be used to fine-tune immune responses during enteric bacterial infections to maintain host protection while reducing immunopathology.

## Results

### Inhibition of Substance P receptor signaling reduces the severity of *C. rodentium* infection

To ascertain the role of SP receptor (TACR1) signaling in the host-response to enteric infection with *C. rodentium*, mice were treated with vehicle or the highly selective receptor antagonist CP96345 (2.5 mg/kg, daily orogastric gavage). Bacterial burden assessed in feces revealed reduced bacterial burden throughout the infection, reaching statistical significance for 10-, 18-, and 21-days post-infection **(Figure 1A)**. In separate cohorts of mice, statistically significant reductions were again observed 10 dpi in fecal pellets **(Figure 1B)**, colonic tissues **(Figure 1C)**, and in the lumen of colonic tissue sections by confocal microscopy **(Figure S1A&B)**. As expected, vehicle-treated *C. rodentium-*infected mice had significantly increased colonic crypt hyperplasia compared to non-infected controls. In contrast, treatment with CP96345 reduced crypt hyperplasia 10 dpi **(Figure 1D&E)**. These morphometric data were supported by significantly reduced numbers of proliferating (Ki67+) epithelial cell (CDH1+) cells in CP96345-treated *C. rodentium*-infected mice compared to vehicle 10 dpi **(Figure 1F&G)**. While increased crypt length and epithelial cell proliferation were still observed at 29 dpi compared to uninfected controls, there was no difference between treatment groups in *C. rodentium* infected mice 29 dpi. We confirmed that *C. rodentium* viability was not reduced by CP96345 at equivalent concentrations used in our *in vivo* studies **(Figure S1C)**. Additionally, *C. rodentium* cultured in DMEM showed no defect in mRNA expression of the LEE transcriptional regulator *ler*, or the regulated genes *espA,* or *espB* in the presence of physiologically relevant concentrations of CP96345 when compared to vehicle controls **(Figure S1D-F)**. These data indicate that CP96345 does not impede the ability of *C. rodentium* to express these virulence genes. We further assessed colonic motility as a possible mechanism for reduced bacterial burden; however, we observed reduced colonic motility in CP96345-treated mice **(Figure S1G)** demonstrating that these reductions in bacterial burden are not simply due to increased colonic motility. Highlighting that our observations are due to reduced TACR1 signaling and not a CP96345-specific effect, we observed reduced *C. rodentium* burden in mice treated with the antagonist SR140333 (1 mg/kg)^26^ delivered by intraperitoneal injection **(Figure S1H&I)**. *C. rodentium* is well-known for increasing oxygen availability as a mechanism aiding bacterial colonization and pathogenesis. Given that we saw significant decreases in bacterial burden and pathology, we investigated if oxygenation was a potential mechanism. We observed equivalent staining of hypoxyprobe in CP96345-treated mice compared to respective controls **(Figure S1J)**. Together these data demonstrate that TACR1 signaling is a critical element of host responses during enteric bacterial infection that reduces bacterial burden and infection-induced pathology.

**Figure 1.**
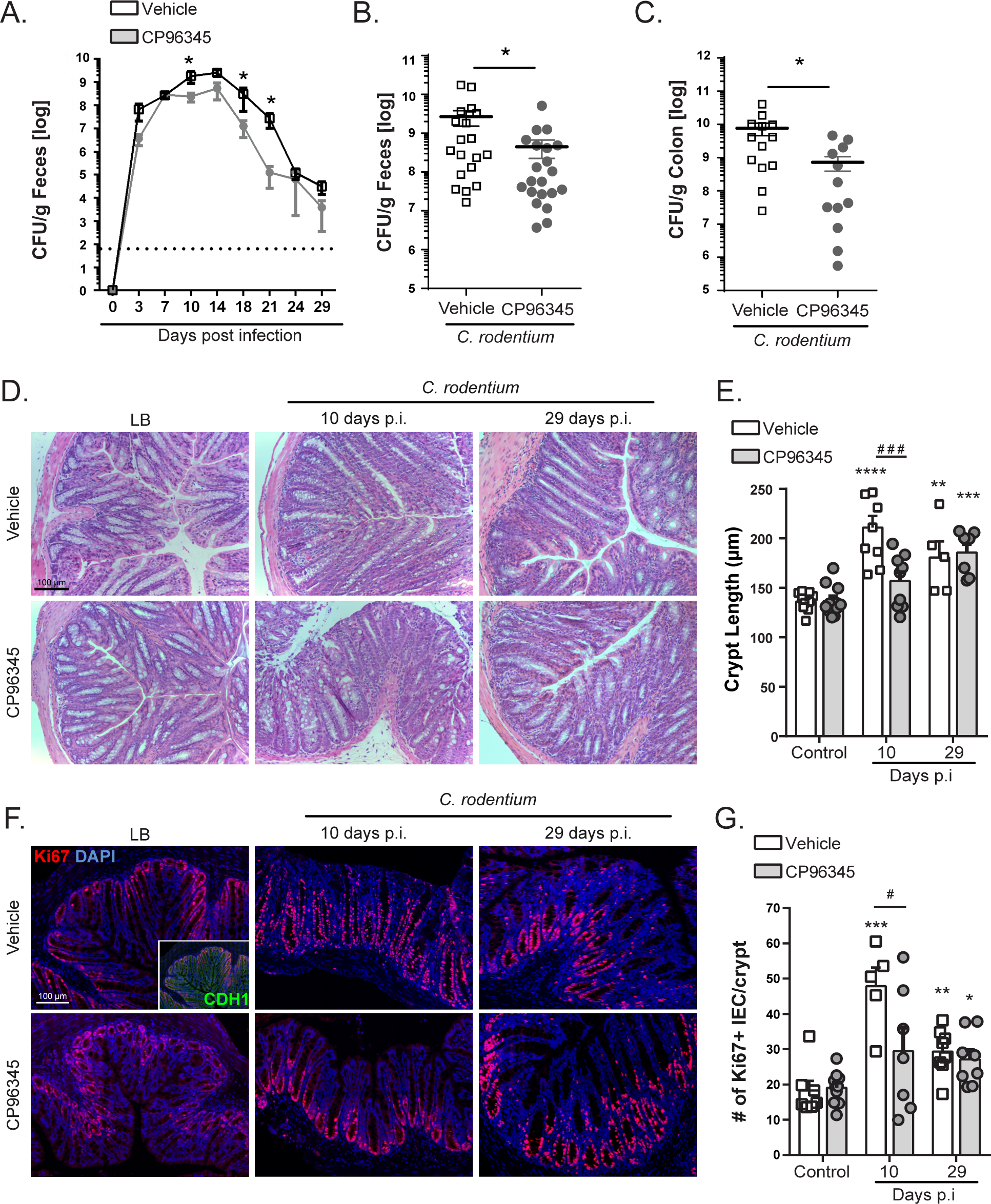
Antagonism of TACR1 signaling reduces *C. rodentium* burden and induced pathology. Administration of the highly selective TACR1 antagonist CP96345 (2.5 mg/kg, orogastric gavage, once daily) significantly reduces fecal *C. rodentium* over the course of infection **(A)**. Reduced fecal **(B)** and colonic adherent **(C)** *C. rodentium* were observed in CP96345 compared to vehicle-treated mice at 10 days post-infection (dpi). Crypt length was measured on H&E-stained histology sections **(D)** and quantified **(E)** from vehicle or CP96345-treated uninfected or infected mice at 10 and 29 dpi. Immunofluorescent staining and confocal imaging were performed to identify **(F)** and enumerate **(G)** proliferating (DAPI+ Ki67+ CDH1+) IEC in uninfected or infected mice treated with vehicle or CP96345. Results are from individual mice, mean ± SEM. For **(A-C)** * = P ≤0.05, ** = P ≤0.01, *** = P ≤0.001. For **(D-G)**, * = P ≤0.05, ** = P ≤0.01, *** = P ≤0.001 compared to uninfected controls of the same treatment and # = P ≤0.05, ## = P ≤0.01, ### = P ≤0.001 compared between treatment groups. Two-way ANOVA with Tukey’s post-hoc test. Scale bar = 100µm. Luria broth (LB), post-infection (p.i.), intestinal epithelial cells (IEC).

### Colonic T-cell recruitment is abrogated by antagonism of the Substance P receptor

Colonic crypt hyperplasia due to *C. rodentium* infection is well established to be driven by IFNγ-producing T-cells^22,24^. With this in mind, we hypothesized that reduced colonic hyperplasia could be due to decreased T-cell recruitment or production of these cytokines. Confocal microscopy revealed that, as expected, *C. rodentium* infection significantly increased the number of T-cells (CD3+ DAPI+) in colonic tissue sections when compared to non-infected mice. However, the number of colonic T-cells at 10 and 29 dpi with *C. rodentium* was significantly reduced in CP96345-treated mice compared to vehicle-treated controls **(Figure 2A&B)**. These results were confirmed using flow cytometry demonstrating reduced numbers of CD3+ T-cells (Live, singlet, CD45+, CD3+) in infected mice treated with CP96345 compared to vehicle 10 dpi **(Figure 2C)**. Using *ex vivo* restimulation and intracellular cytokine staining, we identified that IFNγ and IL-17A, but not IL-22-producing CD4+ T-cells were reduced in infected mice treated with CP96345 compared to vehicle-treated *C. rodentium*-infected controls **(Figure 2D-F)**. Despite these changes in IFNγ and IL-17A, we found no effect of CP96345 on the prevalence of FoxP3+ T-cells in the lamina propria **(Figure 2G)** or in MLN **(Figure S2A)** across treatment groups. These data demonstrate that TACR1 antagonism reduces T-cell recruitment and cytokine production to reduce host-maladaptive immunopathology.

**Figure 2.**
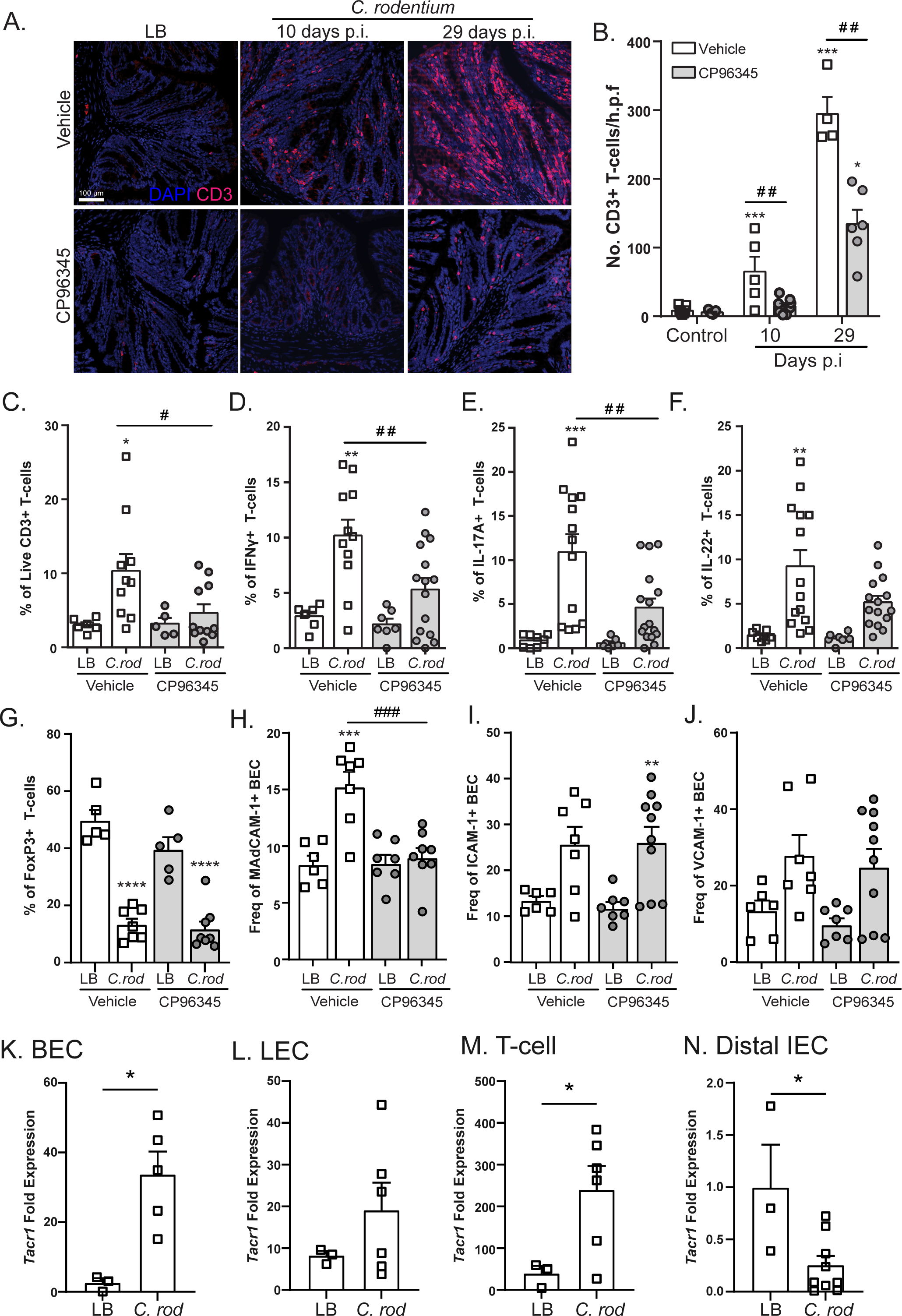
TACR1 signaling is required for recruitment of IFNγ and IL-17A producing T-cells during enteric infection. Colonic recruitment of T-cells (DAPI+ CD3+) in uninfected (LB) or *C. rodentium* infected mice treated with vehicle or CP96345 was assessed in tissues sections obtained 10 and 29 dpi by immunofluorescence and confocal microscopy **(A)** and enumerated **(B)**. LB or *C. rodentium* infected mice treated with vehicle or CP96345 10 dpi were quantified via flow cytometry for T-cell recruitment by frequency of live CD3+ T-cells **(C)** as well as intracellular cytokine staining for CD4+ T-cells that express IFNγ **(D)**, IL-17A **(E)**, IL-22 **(F)** and FoxP3 **(G)**. In a separate cohort, colonic BEC (CD45-, CD31+ gp38-) from uninfected (LB) and *C. rodentium* infected mice treated with vehicle or CP96345 10 dpi were analyzed for their expression of MAdCAM-1 **(H)**, ICAM-1 **(I)**, and VCAM-1 **(J)**. *Tacr1* mRNA expression was assessed in sorted colonic BEC **(K)**, lymphatic endothelial cells (LEC) **(L)**, T-cells **(M)**, and distal IEC **(N)** in naïve (LB) or 10 dpi *C. rodentium* infected WT mice. Each data point represents an individual mouse. Mean ± SEM, * = P ≤0.05, ** = P ≤0.01, *** = P ≤0.001 compared to uninfected controls of the same treatment and # = P ≤0.05, ## = P ≤0.01, ### = P ≤0.001 compared between treatment groups. One-way ANOVA with Tukey’s post-hoc test was used. Scale bar = 100µm. Luria broth (LB), post-infection (p.i.), *Citrobacter rodentium* (*C. rod*).

Reductions in bacterial burden early in the course of disease prior to when CD4+ T-cells are recruited to the colon **(Figure 1A)** led us to assess the ability of TACR1 antagonism to alter innate lymphoid cell (ILC) populations. Focusing on 3 dpi, a time point selected to occur before T-cell responses, we found that CP96345 treatment did not affect the frequency of ILC1, ILC2, or NK cells in the colonic lamina propria regardless of *C. rodentium* infection **(Figure S2B-D)**. Further analysis demonstrated a significantly decreased frequency of natural cytotoxicity receptor (NCR)+ ILC3 and CD4-lymphoid tissue inducer (LTi) cells, with no significant decrease in NCR-ILC3 and CD4+ LTi cells in CP96345-treated compared to vehicle-treated mice **(Figure S2E-H)**. Thus, TACR1 receptor antagonism has marginal effects on specific ILC3 and LTi cell subpopulations.

Given the reduction of T-cell recruitment into the lamina propria, we assessed the expression of cell adhesion molecules on the surface of BEC critical for T-cell extravasation from the vasculature. By flow cytometry, we found a significant increase in frequency of ICAM-1+, VCAM-1+ and MAdCAM-1+ BEC (CD45-, gp38-, CD31+) in *C. rodentium*-infected mice 10 dpi compared to uninfected controls. BECs from infected mice receiving CP96345 upregulated ICAM-1 and VCAM-1 but had significantly reduced cell surface MAdCAM-1 expression compared to infected vehicle-treated controls **(Figure 2H-J)**. Additionally, we found that sorted colonic BEC, lymphatic endothelial cells (LEC: CD45-, gp38+, CD31+), and T-cells all express high levels of TACR1 mRNA relative to a known expressing ganglion. Specifically, BEC and T-cells show significantly increased *Tacr1* expression 10 dpi with *C. rodentium* **(Figure 2K-M)**. Conversely, distal IEC express *Tacr1* but significantly downregulate expression during *C. rodentium* infection **(Figure 2N).** These data suggest that TACR1 antagonism reduced colonic T-cell recruitment by reducing blood endothelial MAdCAM-1 upregulation in a cell-intrinsic manner.

### Substance P receptor signaling attenuates host inflammation during *C. rodentium* infection

With the reduced colonic T-cell recruitment and MAdCAM-1 expression, we assessed if CP96345 abrogated the expression of proinflammatory cytokines during *C. rodentium* infection. In keeping with our flow cytometry data, CP96345-treated *C. rodentium*-infected mice had significantly reduced *Ifnγ* mRNA compared to vehicle-treated infected mice 10 and 29 dpi **(Figure 3A)**. *Il17a* was also significantly decreased by CP96345 treatment at 29 dpi compared to vehicle-treated controls **(Figure 3B)** while *Il22* was not affected by treatment at either timepoint **(Figure 3C)**. Confirming that these results were not specific to CP96345, administration of the TACR1 antagonist SR140333 reduced *Ifnγ* expression at 10 dpi compared to vehicle-treated *C. rodentium*-infected mice **(Figure S3A)**, although SR140333 did not alter *Il17a* or *Il22* expression **(Figure S3B&C)**.

**Figure 3.**
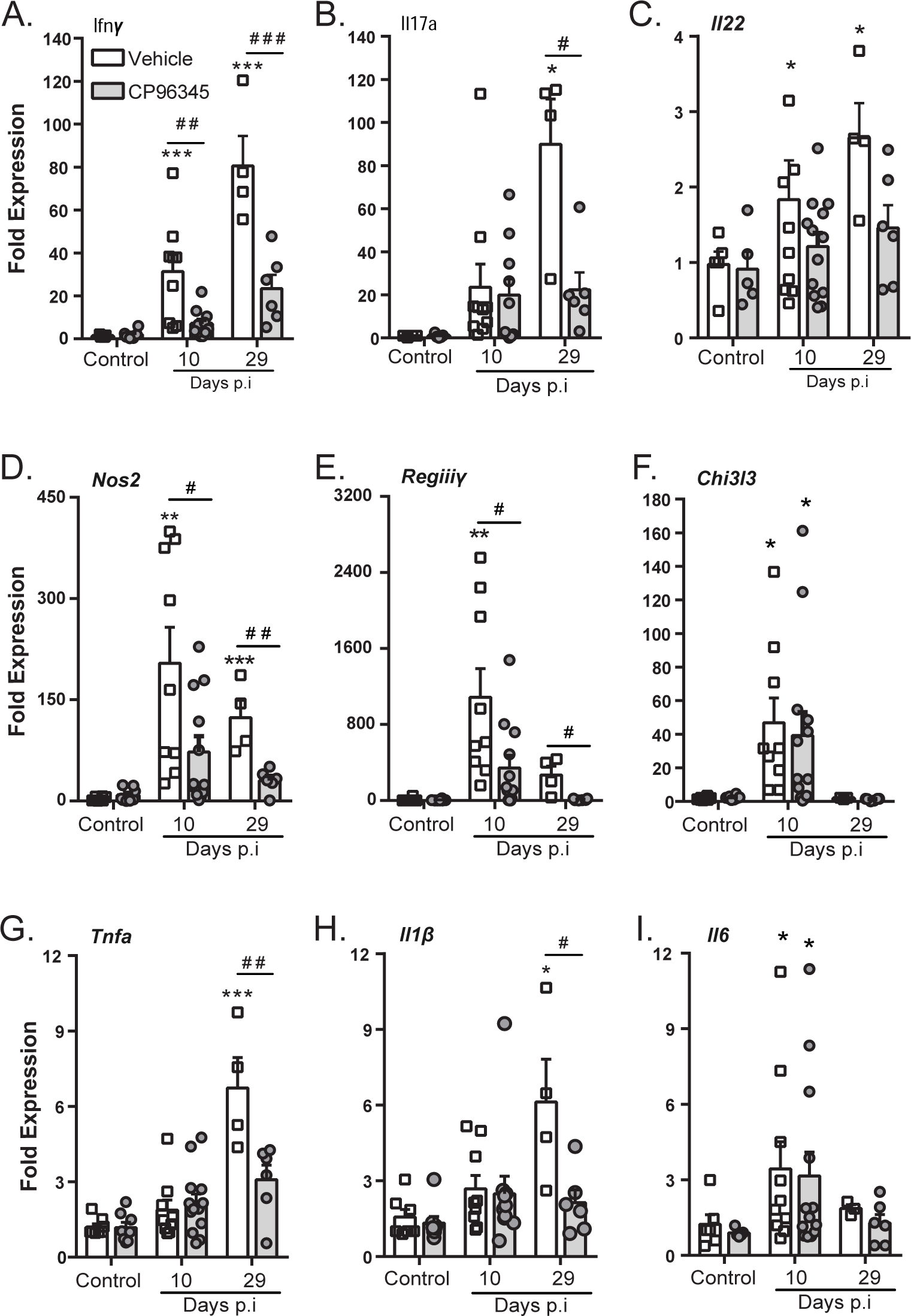
TACR1 blockade attenuates expression of specific pro-inflammatory genes in colonic tissues during *C. rodentium* infection. Colonic tissue from uninfected (LB) and infected mice treated with vehicle or CP96354 were assessed for expression of *Ifnγ* **(A)**, *Il17a* **(B)**, *Il22* **(C)** at 10 and 29 dpi. General inflammatory responses were assayed by measuring colonic expression of *Nos2* **(D)**, *Regiiiу* **(E)**, *Chi3l3* **(F)**, *Tnfa* **(G)**, *Il1β* **(H)**, and *Il6* **(I)**. Results are from individual mice, mean ± SEM, * = P ≤0.05, ** = P ≤0.01, *** = P ≤0.001 compared to uninfected controls of the same treatment and # = P ≤0.05, ## = P ≤0.01, ### = P ≤0.001 compared between treatment groups. Two-way ANOVA with Tukey’s post-hoc test was used. Post-infection (p.i.).

IFNγ is well appreciated to increase the expression of a variety of genes that are critical to host protection, including inducible nitric oxide synthase (iNOS, *Nos2*) and RegIIIγ^24,27^. Infection-induced increases in *Nos2* expression at 10 and 29 dpi were significantly reduced in CP96345-treated mice **(Figure 3D)**. *Regiiiγ* is a critical antimicrobial peptide often upregulated during *C. rodentium* infection in mice. We show similar upregulation of *Regiiiγ* induced by *C. rodentium* infection, but CP96345 treatment significantly reduced *Regiiiγ* expression at 10 and 29 dpi compared to vehicle **(Figure 3E).** With TACR1 antagonism showing modulation of genes related to macrophage function, we investigated *Chi3l3,* the alternatively activated macrophage marker gene, which was induced by *C. rodentium* infection but was not affected by treatment with CP96345 compared to vehicle controls at similar time points **(Figure 3F)**. In support of SP receptor regulation of specific gene expression profiles in response to enteric bacterial infection, *Tnfa* and *Il1β* expression induced by infection were significantly reduced by TACR1 antagonism 29 dpi compared to vehicle-treated and infected mice **(Figure 3G&H)**. Expression of *Il6* induced by *C. rodentium* infection was not impacted by treatment with CP96345 **(Figure 3I)**. These results demonstrate TACR1 signaling during *C. rodentium* infection is a critical modulator of specific host-protective gene expression during inflammation, serving to shape the host response.

### Antagonism of Substance P receptor does not prevent antigen-specific T-cell proliferation *in vivo*

Here we assessed if reduced colonic T-cell recruitment during TACR1 antagonist treatment was due to abrogated dendritic cell migration or function and antigen-specific T-cell proliferation. No significant differences in cDC (CD45+ CD3-B220-CD317-CD11c^hi^ MHCII^hi^) from the mesenteric lymph nodes (MLN) were identified 10 dpi in LB or *C. rodentium* infected mice treated with vehicle or CP96345 **(Figure S4A)**. There was also no significant difference in either protein expression of the costimulatory molecule CD86 by cDC **(Figure S4B)** or in the number of CD11b- or CD11b+ CD103+ migratory cDC **(Figure S4C)**. To assess if TACR1 antagonism reduced the function of DC to present antigen to naïve antigen specific T-cells, Cell Proliferation Dye eFluor450 labelled CD4+ OT-II T-cells were adoptively transferred into WT mice and treated with vehicle or CP96354, then gavaged with ovalbumin or PBS **(Figure S4D)**. TACR1 antagonism did not alter OT-II T-cell proliferation within the MLN measured by proliferation and division index **(Figure S4E&F)**; however, there was a significant increase in the number of OT-II T-cells in MLN in CP96345-treated mice compared to vehicle controls **(Figure S4G)**. Together these data show that TACR1 antagonism neither alters the number and function of cDC, nor the ability to induce antigen specific T-cell proliferation but could reduce lymph node egress.

### T-cell intrinsic Substance P receptor signaling enhances IFNγ production

To decipher the role of T-cell intrinsic TACR1 signaling during *C. rodentium* infection, we used a conditional knockout approach. TACR1 T-cell conditional knockout (Lck.Cre+ TACR1^f/f^) and WT (Lck.Cre-TACR1^f/f^) littermate controls were infected with *C. rodentium* and assessed 10 dpi. Flow cytometry conducted on colonic lamina propria lymphocytes revealed significant increases in T-cell recruitment in both T-cell cKO and WT mice compared to LB treated controls with no difference between genotypes **(Figure 4A)**. In assessing cytokine production by intracellular cytokine staining and flow cytometry, the frequency of IFNγ+ T-cells was significantly reduced in infected TACR1 T-cell cKO compared to WT mice 10 dpi **(Figure 4B)**. Although a slight reduction in IL-17A+ T-cells was also observed in infected TACR1 T-cell cKO compared to WT mice 10 dpi, this was not statistically significant **(Figure 4C).** As expected, *C. rodentium* infection increased IL-22+ colonic T-cells; however, this was not different in WT compared to TACR1 T-cell cKO mice **(Figure 4D)**. Regulatory FoxP3+ CD4+ T-cells were also not different between WT or T-cell TACR1 cKO mice in the lamina propria **(Figure 4E)** and the MLN **(Figure S5A-B)**. These data were confirmed in qPCR conducted on colonic tissue samples, with infection-induced increases in *Ifnγ* expression in WT mice that were significantly reduced in TACR1 T-cell cKO mice 10 dpi **(Figure 4F)**. Expression of *Il17a* and *Il22* showed no difference between *C. rodentium*-infected WT and TACR1 T-cell cKO mice 10 dpi **(Figure 4G&H)**. With the reduced IFNγ production by TACR1 T-cell cKO compared to WT mice during infection, we assessed if T-cells from these mice were able to proliferate and differentiate to Th1 or Th17 T-cells. Using *in vitro* differentiation, we found no defect in TACR1 T-cell cKO proliferation by EdU incorporation, or Th1 or Th17 differentiation indicated by production of IFNγ and IL-17A respectively **(Figure S5C-F)**. Interestingly, we found no significant difference in bacterial burden in these T-cell cKO compared to WT mice **(Figure S5G&H)**. Infection with *C. rodentium* induced crypt hyperplasia, with no difference in crypt length in WT compared to TACR1 T-cell cKO mice 10 dpi. **(Figure 4I&J)**. In keeping with these data, we also saw no difference in proliferating (Ki67+) intestinal epithelial cells across genotypes **(Figure 4K&L).** Together, these data demonstrate that TACR1 cell-intrinsic signaling in T-cells reduces select cytokine responses but is not sufficient to reduce bacterial burden and host pathology.

**Figure 4.**
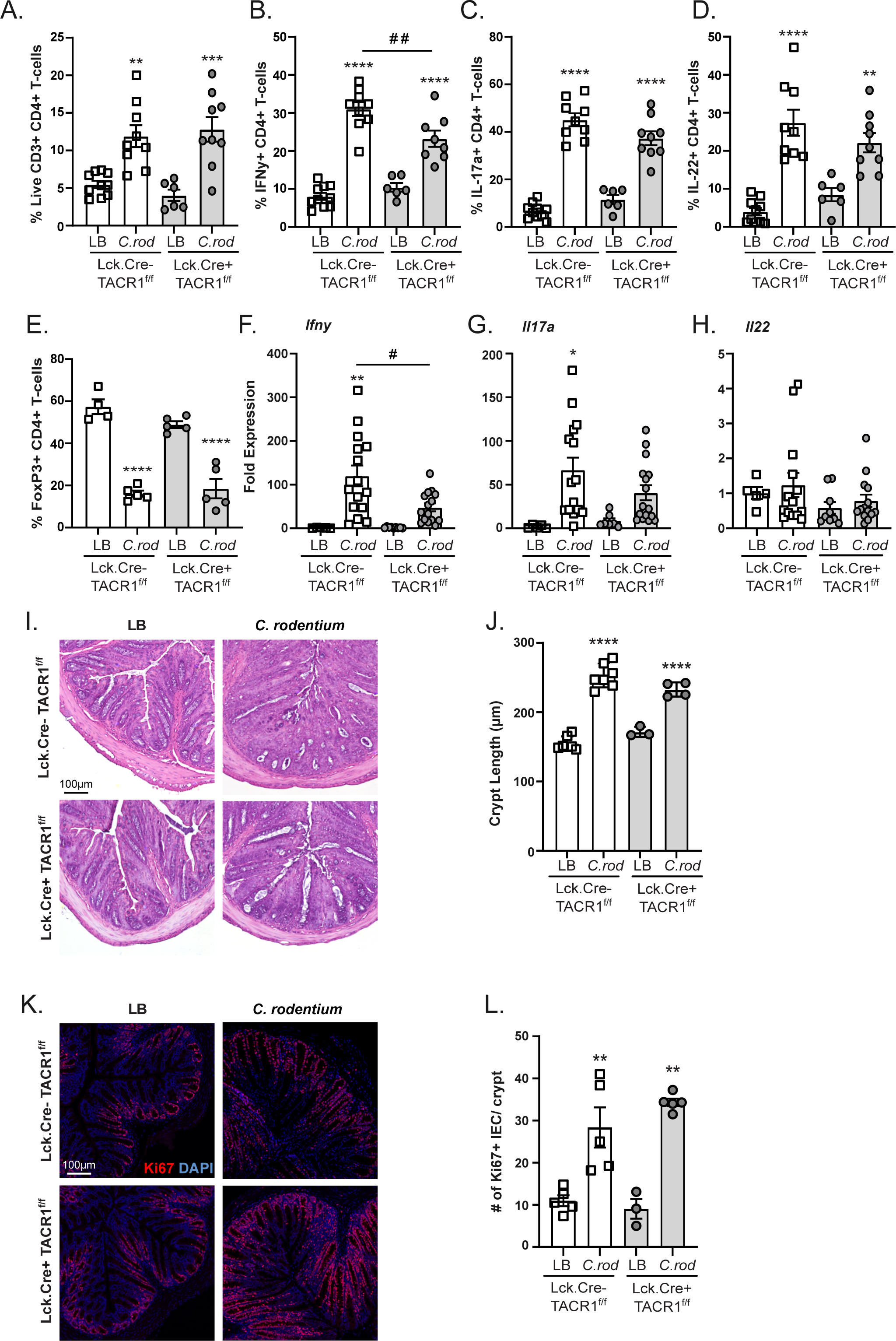
T-cell intrinsic TACR1 signaling impacts IFNγ production during enteric infection. Colonic lamina propria lymphocytes were quantified in Lck.Cre+ TACR1^f/f^ mice and their Lck.Cre-TACR1^f/f^ littermates. **(A)** Frequency of live CD3+ CD4+ T-cells in the colonic lamina propria of *C. rodentium* infected mice 10 dpi. Intracellular cytokine staining of IFNγ **(B)**, IL-17A **(C)**, IL-22 **(D)**, and FoxP3 **(E)** producing CD4+ T-cells 10 days post-*C. rodentium* infection. Colonic tissue from Lck.Cre+ TACR1^f/f^ mice and their Lck.Cre-TACR1^f/f^ littermates was assessed for mRNA expression of *Ifnγ* **(F)**, *Il17a* **(G)**, *Il22* **(H)** at 10 dpi. **(I&J)** Colonic crypt length was measured in H&E-stained sections of LB or 10 dpi *C. rodentium* infected Lck.Cre+ TACR1^f/f^ mice or Lck.Cre-TACR1^f/f^ littermates. DAPI+ Ki67+ CDH1+ IEC were stained **(K)** and enumerated **(L)** via immunofluorescent staining and confocal imaging in uninfected or infected Lck.Cre+ TACR1^f/f^ mice and their Lck.Cre-TACR1^f/f^ littermates 10 dpi. Results are from individual mice, mean ± SEM, * = P ≤0.05, ** = P ≤0.01, *** = P ≤0.001 compared to uninfected controls of the same treatment and # = P ≤0.05, ## = P ≤0.01, ### = P ≤0.001 compared between treatment groups. One-way ANOVA with Tukey’s post-hoc test was used. Scale bar = 100µm. Luria broth (LB), *C. rodentium* (*C. rod*).

## Discussion

Host-protective responses elicited by pathogens in the intestinal tract are tightly controlled and require a diverse array of cell types. Coordination of these responses can occur through many cell-intrinsic and cell-cell communication networks^28^. Neurotransmitters released by the neurons within the intestinal tract have become increasingly recognized to exert not only control over intestinal physiology but also immune function. We have previously demonstrated that ablation of sensory neurons expressing TRPV1 resulted in increased bacterial burden, and reduced T-cell recruitment^8^. This role of TRPV1 in sensory neurons was further demonstrated to elicit host protective recruitment of neutrophils during *C. rodentium* infection^9^. These data suggested that there could be a complex array of signals that emanate from sensory neurons during enteric bacterial infection. SP has long been identified to function as a neurotransmitter critical in mediating neurogenic inflammation, intestinal physiology and host-protective immune responses. The localized release of SP serves not only as a nociceptive neurotransmitter but also acts through the SP receptor on endothelial cells to increase vascular permeability, adhesion molecule expression, and vessel dilation (smooth muscle)^1^. Despite these roles and our prior studies demonstrating the host protective role of TRPV1+ sensory nociceptors^8,9^, the specific function of SP during enteric bacterial infection was not well established. Our experiments indicate that SP receptor signaling modulates immune responses to *C. rodentium* infection that can be host maladaptive. Here we demonstrate that SP receptor antagonism reduces bacterial burden and colonic crypt hyperplasia. This reduced bacterial burden was not simply due to bactericidal or bacteriostatic effects and could be replicated in experiments using a different SP receptor antagonist administered by i.p. injection. Additionally, we observed significantly decreased *Tnfα* and *Il1β* at the end of the infection, suggesting that SP receptor antagonism reduced in bacterial burden and pathology led to lasting effects in the host after the pathogen is cleared. These results are in keeping with recent findings where SP KO mice or, blockade of SP receptor signaling reduced intestinal inflammation induced by *Clostridium difficile* infection^29^. Together these data demonstrate that SP receptor signaling is a critical determinant of bacterial burden during *C. rodentium* infection and suggest that TACR1 signaling can act in a host-deleterious manner.

To discern how SP receptor antagonism may reduce *C. rodentium* bacterial burden and host pathology, we considered that treatment may limit the availability of host-derived factors required for maximal colonization by the pathogen^30–32^. Proliferation of *C. rodentium* in the colonic lumen requires inflammation to increase oxygen availability in an otherwise hypoxic environment^33^. However, there was no difference in colonic epithelial oxygenation in control or infected mice treated with SP receptor antagonist compared to vehicle controls. These data indicate that blocking SP receptor signaling reduces *C. rodentium* bacterial burden and inflammation, independent of increased colonic oxygenation.

Intestinal pathology during *C. rodentium* infection is not only due to the presence of this attaching and effacing pathogen but is also a consequence of the host’s immune response to the infection. Prior elegant studies demonstrated that increased IEC proliferation that causes crypt hyperplasia is due to the recruitment of CD4+ T-cells producing IFNγ and IL-22^22,23^. This IFNγ-dependent immunopathology can however be fatal in the absence of macrophage and CD11b+ CD103-dendritic cells that produce IL-23^34^. Here we demonstrate that SP receptor antagonism significantly reduced IEC proliferation (Ki67+ CDH1+), and consequently, infection-induced crypt hyperplasia at 10 dpi but not 29 dpi compared to infected vehicle-treated mice. This appeared to not be driven by IEC intrinsic TACR1 signaling as IEC expression of *Tacr1* was low and further downregulated during *C. rodentium* infection, suggesting that the crypt hyperplasia is driven by cell-cell interactions instead.

IFNγ signaling is well established to regulate the response to *C. rodentium* by acting on a variety of diverse cell types. As the primary interface between the host and bacterial pathogen, proteomic analysis on colonic IEC identified infection-induced increases in proteins regulated by IFNγ such as NOS2, RegIIIγ, and major histocompatibility complex II (MHCII). Although IFNγ deficiency did not increase bacterial burden at the height of the disease, IFNγ KO mice failed to increase MHCII on colonic epithelial cells and clear the infection. This requirement for MHCII expression by colonic IEC in protecting from *C. rodentium* infection was however discounted as IFNγ KO mice produced anti-*C. rodentium* antibodies and were protected from re-infection^24^. Subsequent analysis confirmed that IFNγ signaling in IEC does not affect bacterial burden but instead is critical in driving intraepithelial lymphocytes to restrain infection-induced inflammation^35^. Our data with SP receptor antagonists reducing colonic IFNγ and recruitment of IFNγ+ CD4+ T-cells suggests that this is one facet of the immune response capable of reducing intestinal inflammation. Expression of MHCII on mid-distal IEC has also been suggested to present antigen to IL-22+ CD4+ T-cells, sustaining cytokine release and aiding in the removal of susceptible IEC. Blockade of SP receptor signaling or conditional ablation of SP receptor on T-cells reduced but did not ablate IFNγ production, suggesting that a fine balance of inflammation can be achieved, allowing for host protection without induction of immunopathology. Together these data indicate that host response to intestinal inflammation can be tuned through a variety of mechanisms including SP receptor signaling.

To understand the role of SP receptor signaling in T-cell recruitment and given the known ability of SP to increase adhesion molecule expression that are critical to immune cell homing during intestinal inflammation^36,37^, we performed flow cytometry to assess the surface expression of these proteins on colonic BEC. In keeping with our prior results, *C. rodentium* infection increased adhesion molecule mRNA expression^8^ and protein on the surface of colonic BEC^9^. Compared to vehicle-treated *C. rodentium*-infected mice, MAdCAM-1 expression was significantly reduced on the endothelial cell surface of SP receptor antagonist-treated mice. Our data further suggests this may be driven by SP receptor signaling on BEC intrinsically as BEC express *Tacr1* mRNA which is further upregulated during *C. rodentium* infection. These results suggest that SP receptor signaling enhances colonic T-cell recruitment by increasing MAdCAM-1 on the surface of endothelial cells. Although SP has been reported to increase adhesion molecule expression such as ICAM-1 and VCAM-1^28,36,38^, SP receptor antagonism did not reduce these cellular adhesion molecules induced by *C. rodentium* infection. It is critical to note that the SP-induced ICAM-1 and VCAM-1 expression were conducted on human dermal microvascular endothelial cells, or endothelial cell lines *in vitro*^36,37^. Mouse endothelial cell lines have also been shown not to increase ICAM-1 or VCAM-1 expression in response to SP^39^. These differences could suggest that SP receptor signaling in intestinal BEC may elicit unique responses or that there are species-specific differences in SP receptor expression or signaling. Expression of MAdCAM-1 by BEC in the intestine has long been known as critical to the recruitment of mucosal homing T-cells through interacting with T-cells bearing α4β7 on their surface^40–43^. MAdCAM-1 expression is increased in models of colitis, and colitis severity can be reduced through blockade or neutralization of α4β7. These findings therefore indicate that SP receptor signaling blockade can reduce infection-induced immunopathology by attenuating the expression of key adhesion molecules that recruit pathogenic T-cells into the colonic lamina propria.

Using a novel SP receptor T-cell cKO mouse, we further uncovered a unique role of T-cell intrinsic signaling in *C. rodentium*-induced pathogenesis. Infected SP receptor T-cell cKO mice had reduced colonic IFNγ expression and numbers of CD4+ IFNγ producing T-cells. Additionally, we found that lamina propria recruited T-cells upregulate *Tacr1* mRNA during *C. rodentium* infection, suggesting that the enhanced IFNγ expression may be driven by local tissue signaling. However, recruitment of T-cells to the infected colon is not SP receptor T-cell intrinsic, as *C. rodentium* infection induced equivalent T-cell recruitment in WT and cKO mice. These data are in keeping with previous publications demonstrating selective SP receptor antagonists significantly reduced antigen-specific IFNγ release upon T-cell restimulation from *Schistosoma mansoni* infected mice^44^, and that SP enhances IFNγ^45^. Critically, we found that CD4+ T-cells deficient in the SP receptor were still able to differentiate towards Th1 and Th17 T-cells *in vitro*, indicating that SP can enhance but is not a requirement for production of these cells. With these data and a prior report of SP receptor enhancing T-cell receptor signaling^25^, we assessed the effect of SP receptor antagonism on antigen-specific T-cell proliferation *in vivo*. Our data demonstrates that antagonism of the SP receptor did not impact OT-II T-cell proliferation in the MLN in response to orally administered ovalbumin, indicating that DC remains capable of antigen uptake, processing, homing to MLN, and presentation to T-cells that remain responsive within these draining LN. Interestingly, we found a significant increase in the number of OT-II cells found in the MLN in mice treated with the SP receptor antagonist compared to vehicle control. This is in keeping with literature demonstrating that injection of SP into the lymph node increased lymph flow and trafficking of immune cells^46^. These experiments demonstrate that the lack of recruited T-cells, and consequently reduced immunopathology is not due to migratory DC deficits, or inability to activate T-cells, but instead suggests defective T-cell homing or emigration.

Together our data highlight that modulation of SP receptor signaling could be used to fine-tune immune responses in the intestine. Antagonism of the SP receptor allows for the development of host protection while attenuating immunopathology induced by enteric bacterial infection. Modulation of immune responses through this signaling axis may prove amenable as our data adds to the pre-existing literature indicating that SP enhances but is not required for immune function, allowing for antagonism without fear of inducing complete immunosuppression.

## Materials and Methods

### Mice

Female and male 6–8-week-old C57BL/6J, and Lck.Cre+ and Lck.Cre-TACR1^f/f^ mice on a C57BL/6J background were used in these studies. Mice were purchased from Jackson Laboratories (Bar Harbor, ME). Cryogenically frozen TACR1 embryos (EM:08399) were provided Consiglio Nazionale delle Ricerche as part of INFRAFRONTIER/EMMA (www.infrafrontier.eu, PMID: 25414328). Mice were re-derived at the Mouse Biology Program (UC Davis). Recovered mice were crossed to FLP mice on a C57BL/6J background to remove a Neo cassette and produce the conditional ready flanked by LoxP site line. These mice were further crossed to C57BL/6J mice, and progeny carrying the allele of interest were subsequently bred with Lck.Cre+ mice. All procedures and protocols were approved by the Institutional Animal Care and Use at UC Davis.

### *Citrobacter rodentium* infection and enumeration of bacteria in feces and colon

*C. rodentium*, strain DBS100, was kindly provided by Dr. Andreas Baumler. Bacteria were grown on MacConkey agar plates overnight at 37°C, followed by an overnight culture in Luria broth (LB) at 37°C without shaking. Mice were gavaged with LB or C. rodentium (10^8^ CFU (colony-forming units)).

Quantification of *C. rodentium* was performed with feces or 1 cm of distal colonic tissue placed into a pre-weighed tube and weighed to allow for determination of sample weight. Samples were homogenized using a stainless-steel bead (Qiagen, Germantown MD) in a tissue lyser (Qiagen), followed by serial dilution and plating on MacConkey agar. Colonies were counted after 16 h of incubation at 37°C, and CFU was calculated per gram of sample.

### TACR1 Antagonist Treatment

Cohorts of mice received vehicle (DMSO diluted in PBS) or the selective substance P receptor antagonist CP96345 2.5 mg/kg by orogastric gavage (Tocris, Minneapolis, MN) from the day of infection until the end of the experiment. Additional cohorts of mice received vehicle or the selective SP receptor antagonist SR140333 (Tocris, Minneapolis, MN) by intraperitoneal injection 1 mg/kg every 2 days for the duration of the experiment.

### Gastrointestinal motility

Motility was measured in vehicle- and antagonist-treated mice after the second dose of the drug, by assaying the number of fecal pellets, in a 20-minute period.

### RNA isolation and Quantitative PCR

Tissues removed during necropsy were placed immediately into TRIzol and frozen until RNA extraction was performed. Samples were thawed, and a stainless-steel bead added to allow for tissue disruption and homogenization in a tissue lyser (Qiagen). RNA extraction was completed as instructed by manufacture (Invitrogen, Carlsbad CA), with quantification by NanoDrop. Synthesis of cDNA was performed using an iScript kit with 1 μg of template RNA, following manufacture instructions (Biorad, CA). This cDNA was used to evaluate target gene expression by quantitative real-time PCR with SYBR green incorporation using primer pairs from Primerbank **(Table S1)**.

**Table S1.**
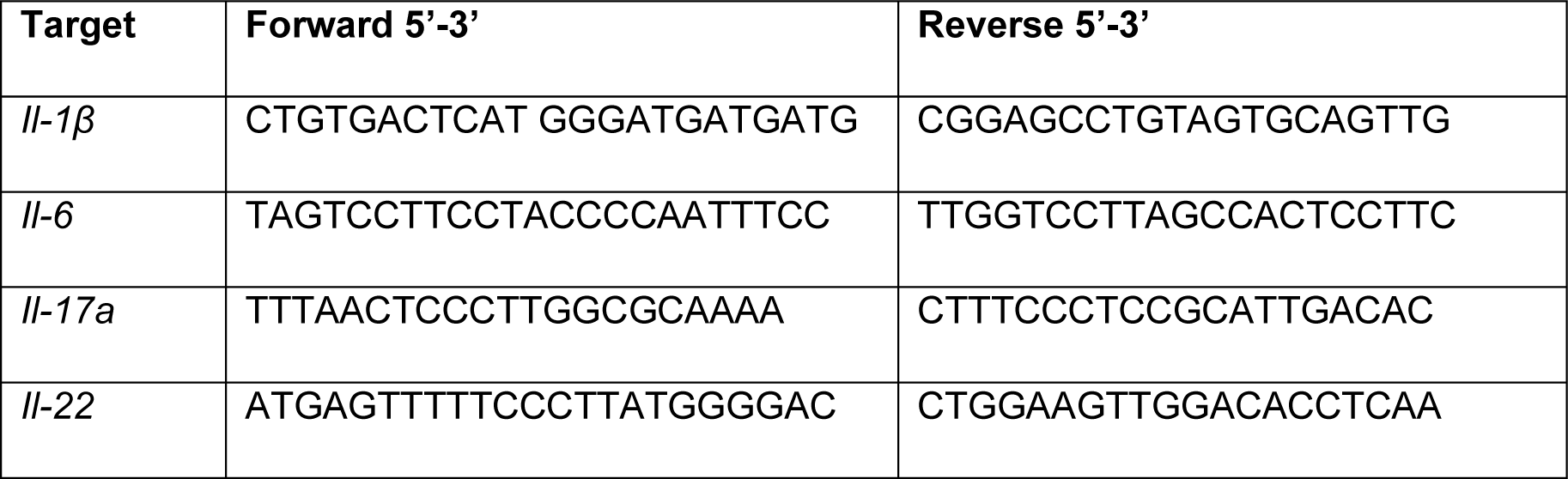
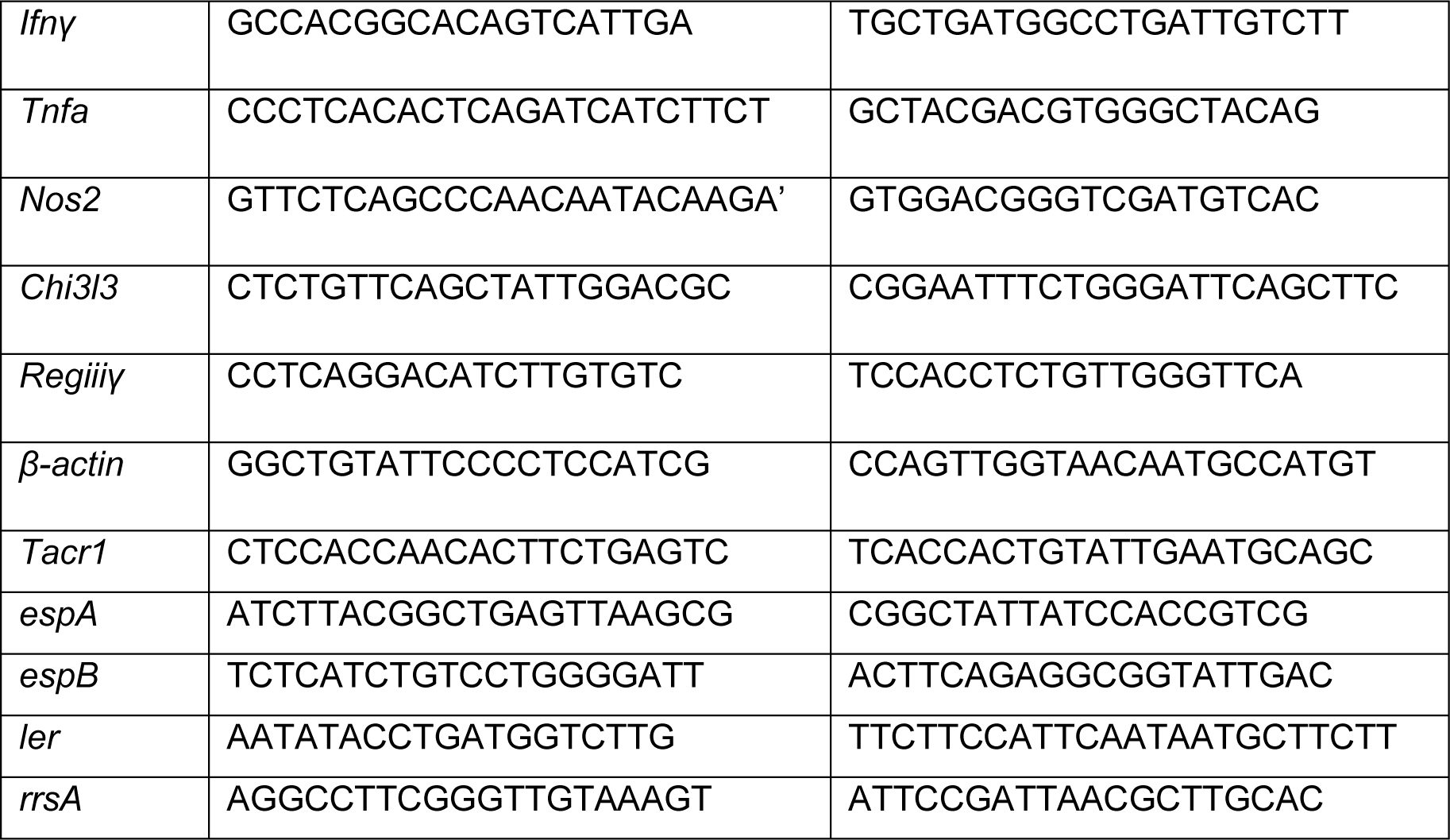
List of primers used for qPCR.

### Histology

Tissues dissected at necropsy were fixed in 10% normal buffered formalin, processed and embedded in Paraffin according to standard protocols. Colonic tissues were embedded on end to allow for cross sections to be obtained. Paraffin embedded samples were sectioned by microtome to produce 6 μm thick sections on slides for histopathology or confocal microscopy. Sections for histopathology were deparaffinized, rehydrated and stained with hematoxylin and eosin following standard protocols. Colonic epithelial cell hyperplasia was assessed by bright field microscopy to allow for measurements of crypt length in at least 15 well-orientated crypts per sample with FIJI (Fiji is just image J, NIH).

### Confocal microscopy

Tissue samples for confocal microscopy were prepared as previously described^9^. In brief, after slides with 6 μm thick sections were de-paraffinized and rehydrated, antigen retrieval was performed in citrate buffer (10 mM, pH 6.0, 30 min., 95°C). After blocking in 5% BSA and normal donkey or goat serum (1 h RT), samples were incubated in primary antibody overnight (16 h, 4°C). Primary and secondary antibodies used are detailed in **Table S2**. Slides were washed extensively (3 x 5 mins) in TBS-tween20 and incubated in appropriately labeled secondary antibodies (Invitrogen Carlsbad, CA) for 1 h at RT, washed, counter-stained with DAPI (1:5000 TBS-tritonX100 0.1% v/v), washed extensively and mounted in Prolong gold (Invitrogen Carlsbad, CA). Staining using anti-mouse CDH1 (E-cadherin) was revealed using a mouse-on-mouse kit according to manufacturer’s instructions (Vector laboratories, Burlingame, CA). Slides were imaged on a Leica SP8 STED 3X confocal microscope with a 40X 1.3 NA objective, with optimal excitation provided by a white-light laser, or a 405 nm laser for DAPI, and appropriate emission collected for each fluorophore. Areas larger than the field-of-view of the objective were acquired using a tiling approach, whereby adjacent images were acquired with a 10% overlap. Files were processed by Imaris Stitcher (Oxford Instruments, UK) to recreate a composite image of the sample. Confocal data analysis was performed by importing Leica image format files into Imaris Stitcher (v9.0, Oxford Instruments) allowing for fusion of overlapping fields-of-view together. Expression of Ki67 in IEC was determined by creating a mask in Imaris (Oxford Instruments) of IEC defined by DAPI+ CDH1+ cells. This masked region was then interrogated for the number of Ki67+ cells. Total Ki67+ cells were divided by the number of crypts defined within the region.

**Table S2.**
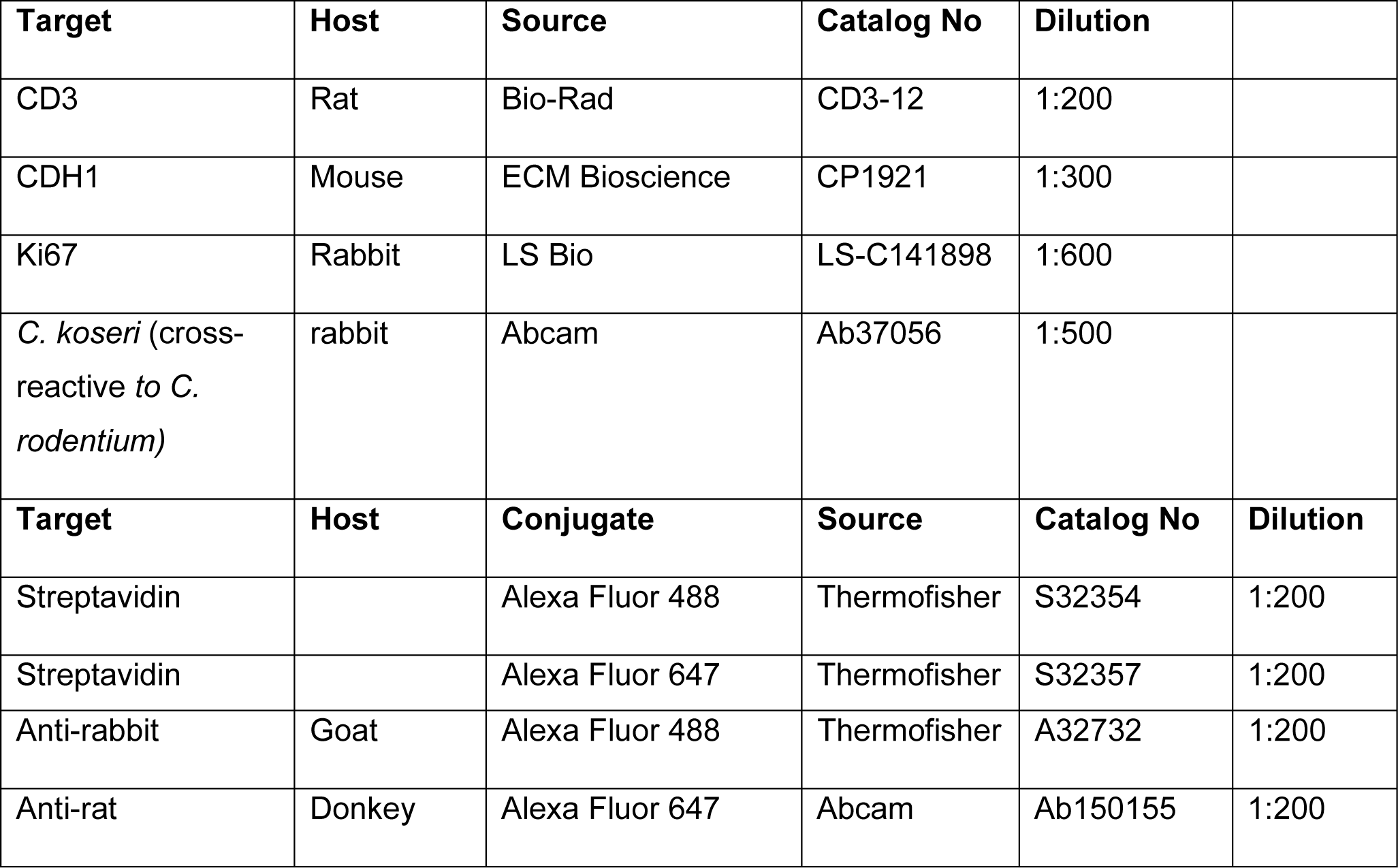
Antibodies used for confocal.

### Colonic hypoxyprobe imaging

To determine the extent of colonic oxygenation, Hypoxyprobe Kit (Cat# HP1-XXX Hypoxyprobe Inc. Burlington, MA) was used. The manufacturer’s protocol was followed. Briefly, WT mice were infected with *C. rodentium* or LB control and received CP96345 or vehicle as described above. After 10 dpi, mice received 2 mg of pimonidazole intraperitoneally 30 minutes before euthanasia, followed by removal and fixation of the colon. Tissues were processed as described above, and Paraffin-embedded sections were stained with mouse anti-pimonidazole (1:50 v/v dilution) followed by detection with streptavidin Alexa Fluor 647, and DAPI to reveal nuclei. Immunofluorescence was imaged as described above and files processed by Imaris (Oxford Instruments).

### Viability of *C. rodentium*

The viability of *C. rodentium* in the presence of vehicle (DMSO) or CP96345 was assessed by the addition of either compound to the bacterial culture. A single colony of *C. rodentium* was cultured overnight in LB at 37°C statically. The bacterial culture was diluted 1:10 in fresh LB with increasing concentrations of CP96345 or DMSO and incubated for 2.5 h at 37°C statically. Culture samples were serially diluted and plated on MacConkey agar plates for overnight incubation before CFU was enumerated.

### Assessment of *C. rodentium* LEE pathogenicity gene expression

A single colony of *C. rodentium* is cultured in LB overnight at 37°C statically. Bacteria is then diluted 1:20 into 10 mL DMEM (Cat# 11995-065 ThermoFisher Waltham, MA) or LB with increasing concentrations of CP96345 or DMSO and cultured at 37°C for 3 hours 200 RPM. Bacteria is pelleted by centrifugation and resuspended in TRIzol reagent. RNA is extracted as described in manufacturer protocol and cDNA generated with iScript reverse transcriptase as described above. Primers for *espA*, *espB*, and *ler* were normalized to the housekeeping gene *rrsA* **(Table S1)**. Expression was compared to the control grown in DMEM with equivalent concentrations of DMSO, whereby a Fold Expression of 1 means there is no difference between CP96345 or its DMSO control.

### T-cell enrichment

Inguinal, mesenteric lymph nodes, and spleen were sterilely excised from non-treated Lck.Cre+ TACR1^f/f^ and Lck.Cre-TACR1^f/f^ mice and placed on a 100 μm filter and dissociated using the plunger of a syringe followed by several washing steps with stain buffer (1X PBS + 2% FBS). Single cell suspensions were treated with Ack lysis solution (5 min RT), before being washed. Cells were then incubated with Fc block (anti-CD16/32, 10 µg/ml, Tonbo Biosciences, San Diego, CA) for 15 minutes on ice. Antibody cocktail was added to surface stain cells for 30 minutes on ice, followed by extensive washing in stain buffer. The following biotinylated antibodies were added to this cocktail at 1:50 dilution: anti-CD161 (clone# PK136, Ref# 30-5941), anti-CD11c (clone# N418, Ref# 30-0114), anti-Ly6G (Clone# RB6-8C5, Ref# 30-5931), anti-TER119 (Clone# TER-119, Ref# 30-5921), anti-CD11b (Clone# M1/70, Ref# 30-0112), and anti-B220 (Clone# RA3-6B2, Ref# 30-0452) from Tonbo Biosciences. Cells were then washed and incubated in magnetic streptavidin beads (Cat# 557812, BD Biosciences, Franklin Lakes NJ) according to the manufacturer’s protocol. Cells were placed in BD IMAG Cell Separation Magnet (Cat# 552311 BD Biosciences, Franklin Lakes NJ) for 8 minutes and the negative fraction was taken and placed into a fresh tube on the IMAG. This process was repeated for two additional magnetic enrichment steps to reach a T-cell purity of >90% confirmed by flow cytometry.

### In vitro T-cell differentiation and ELISA

Negatively enriched CD4+ T-cells were plated on untreated round bottom 96-well plates at 25,000 cells/ well. Wells were pre-treated with 2 µg/mL anti-CD3 (Cat# 40-0032 Tonbo Biosciences San Diego, CA) in PBS for 2 h 37°C 5% CO_2_. Cells were cultured with or without 0.5 µg/mL anti-CD28 (Cat# 40-0281 Tonbo Biosciences San Diego, CA) and in Th1 or Th17 skewing culture conditions. Th1 consisted of 1 µg/mL anti-murine IL-4 (Cat# 16-7041 Invitrogen Carlsbad, CA), 5 ng/mL murine IL-2 (Cat# 212-12 Peprotech Cranbury, NJ), and 10 ng/mL murine IL-12p40 (Cat# 210-12 Peprotech Cranbury, NJ). Th17 consisted of 1 µg/mL anti-murine IFNγ (Cat# 16-7312 Invitrogen Carlsbad, CA), anti-murine IL-4 (Cat# 16-7041 Invitrogen Carlsbad, CA), 1 µg/mL anti-murine IL-2 (Cat# 14-7022 Invitrogen Carlsbad, CA), 20 ng/mL murine IL-6 (Cat# 216-16 Peprotech Cranbury, NJ), and 1 ng/mL murine TGF-β1 (Cat# 7666-MB R&D Systems Minneapolis, MN). Cells were incubated 37°C 5% CO_2_ for 96 h, then stimulated with 1X Cell Stimulation Cocktail (phorbol 12-myristate 13-acetate) (eBioscience, San Diego CA) for 4 h. Supernatant was frozen at −20°C until loaded into an ELISA. Untreated 96 flat bottom plates were coated overnight at 4°C with capture antibody for IFNγ or IL-17A from Invitrogen kits (Cat# 88-7314-22 and Cat# 88-7371-88 respectively from Invitrogen, Carlsbad, CA) and manufacturer protocols were followed. Plates were read on a BioTek Synergy HTX Multi-mode Plate Reader using Gen5 application by the 450nm and 570nm wavelengths. 450 nm wavelength was subtracted from the 570 nm wavelength and the standard curve was determined using 4-paramter logistic fit.

### *In vitro* T-cell proliferation analysis

Splenic, mesenteric and inguinal lymph node negatively selected CD4+ T-cells from Lck.Cre+ TACR1^f/f^ mice and Lck.Cre-TACR1^f/f^ littermates were cultured in 96-well round bottom plates coated with 2 µg/mL anti-CD3 and with or without 0.5 µg/mL anti-CD28 as described above. Cells were cultured for 48 or 72 h before adding 10 µM EdU into culture media. Detection of incorporated EdU was performed according to manufacturer’s instructions (Click-IT EdU AF647 Cat# C10340 Invitrogen, Carlsbad, CA) by flow cytometry.

### Isolation of cells from the colon

To reduce colonic tissue to a single cell suspension, lamina propria dissociation kit (Miltenyi Biotec, Gaithersburg, MD) was used with a gentleMACS tissue dissociator. The manufacturer’s protocol was followed. In brief, colons were excised and opened longitudinally, then cut into 1 cm long segments. Epithelial cells removed by gentle agitation (200 RPM) in HBSS supplemented with 10 mM HEPES, 5 mM EDTA and 5% FBS and saved for RNA extraction in TRIzol reagent. Tissue was digested using MACS digestion enzyme mix as described in manufacturer’s protocol and reduced to a single cell suspension by the gentleMACS device. Cell suspension was passed through a 100 µm strainer, washed extensively, and subjected to staining.

### Flow cytometry

A standard flow cytometry staining protocol was followed. Cells were counted manually by hemocytometer with trypan blue exclusion and single cell suspensions incubated with Fc block (anti-CD16/32, 10 µg/ml, Tonbo Biosciences, San Diego, CA) for 25 minutes on ice before incubation with antibody cocktail **(Table S3)** for 30 minutes on ice. Viability was determined using live/dead aqua according to manufacturer’s instructions (ThermoFisher, Waltham MA). Cells were fixed using BD Cytofix (BD Biosciences, Franklin Lakes NJ) for 25 minutes on ice. All flow cytometry data was acquired on a LSRII (BD Biosciences, Franklin Lakes NJ) using DIVA software, with analysis using FlowJo (Becton Dickinson, Eugene OR).

**Table S3.** Flow cytometry antibodies.

| Antibody Target | Manufacture & Clone | Catalog number |
| --- | --- | --- |
| CD3 | Tonbo Biosciences 145-2C11 | 20-0031 |
| CD4 | BD Biosciences RM4-5 | 553047 |
| CD45 | Invitrogen 30-F11 | 48-0451-82 |
| IFNγ | Invitrogen XMG1.2 | 25-7311-82 |
| IL-17A | BD Biosciences TC11-18H10 | 561020 |
| IL-22 | Invitrogen 1HBPWSR | 46-7221-82 |
| CD31 | Biolegend MEC13.3 | 102533 |
| gp38 | Biolegend 8.1.1 | 127405 |
| ICAM-1 | Biolegend YN1/1.7.4 | 116114 |
| VCAM-1 | Biolegend 429(MVCAM.A) | 105719 |
| MAdCAM-1 | Biolegend MECA-367 | 120710 |
| Fixable Live/Dead Aqua | ThermoFisher | L34957 |
| Vα2 | Invitrogen B20.1 | 12-5812 |
| Vβ5 | Invitrogen MR9-4 | 17-5796 |
| CD11c | Invitrogen N418 | 25-0114 |
| MHCII | Invitrogen M5/114.15.2 | 56-5321 |
| CD103 | BD Biosciences M290 | 562772 |
| CD11b | Invitrogen M1/70 | 45-0112 |
| CD86 | BD Biosciences GL1 | 11-0862 |
| NKp46 | Invitrogen 29A1.4 | 25-3351 |
| RORγT | Invitrogen AFKJS-9 | 12-6988 |
| KLRG1 | Invitrogen 2F1 | 47-5893 |
| Eomes | Invitrogen Dan11mag | 61-4875 |
| CD127 | Invitrogen A7R34 | 15-1271 |
| CCR6 | R&D Systems 140706 | FAB590A |
| CD19 | BD Biosciences ID3 | 561740 |
| Gr-1 | Invitrogen RB6-8C5 | 11-5931 |
| F4/80 | Invitrogen BM8 | 53-4801 |
| CD5 | BD Biosciences Ly-1 | 553020 |
| TCR $\beta$ | BD Biosciences H57-597 | 553170 |
| TCR $\gamma\delta$ | BD Biosciences GL3 | 561996 |
| FoxP3 | Invitrogen FJK-16s | 25-5773 |
| B220 | eBioscience RA3-6B2 | 12-0452 |
| CD317 | eBioscience BST2 | 12-3172 |

### Intracellular staining

Prior to surface staining, colonic single cell suspension was incubated in RPMI media with 10% FBS, 1% Penicillin/ Streptomycin, 2 mM L-glutamine, and BD GolgiPlug (1:500, Cat# 555029, BD Biosciences, Franklin Lakes NJ) for 4 hours at 37°C 5% CO_2_ and stimulated with 1X Cell Stimulation Cocktail (phorbol 12-myristate 13-acetate) (eBioscience, San Diego CA). Following surface staining, cells were fixed and permeabilized using a BD Cytofix/ Cytoperm Fixation/ Permeabilization Solution kit (Cat# 554714 BD Biosciences, Franklin Lakes NJ) followed by intracellular staining with anti-IFNγ, anti-IL-17a, and anti-IL-22 **(Table S3)** in 1X BD Perm/ Wash Buffer (Cat# 554723 BD Biosciences, Franklin Lakes NJ) for 1 h. For FoxP3 staining, cells were fixed and permeabilized using FoxP3 Transcription Factor Staining Kit (Cat# 00-5523 Thermofisher Waltham, MA) for 1 h at 4°C and then stained with anti-FoxP3 for 1 h at room temperature. Cells were then extensively washed and analyzed on a LSRII (BD Biosciences, Franklin Lakes NJ) using DIVA software.

### TACR1 mRNA expression determination

Naïve control (LB) and 10 dpi *C. rodentium* infected WT mice were euthanized, and colons excised. A single cell suspension was achieved as described above and stained with anti-CD3, anti-CD45, anti-gp38, anti-CD31, and Live/ Dead Fixable Aqua. Blood endothelial (BEC: CD45-, CD31+, gp38-), lymphatic endothelial (LEC: CD45-, CD31+, gp38+), and T-cells (CD45+ CD3+) were sorted into separate tubes using an Astrios Cell Sorter (Beckman Coulter Brea, CA). Distal IEC were removed as described above, washed and stored in TRIzol. RNA was extracted and processed for qPCR analysis using the Takara CellAmp Direct TB Green RT-qPCR Kit (Cat# 3735A) according to the manufacturer’s protocol.

### Antigen specific T-cell proliferation *in vivo*

C57BL/6J mice were pre-treated with 2.5 mg/kg CP96345 or vehicle (DMSO:PBS) for 3 days followed by another 4 days of treatment daily. After 3 days of pre-treatment, negatively purified splenic and lymph node CD4+ T-cells from OT-II mice were adoptively transferred to these WT pre-treated mice via retro-orbital injection of 10^7^ cells. OT-II cells were negatively purified as described in T-cell enrichment but underwent an additional incubation with Cell Proliferation dye eFluor450 (Cat# 65-0842-85 eBioscience, San Diego, CA) following the manufacturer’s protocol. One day later, 100 mg of ovalbumin in 200 μL PBS was administered intragastrically (i.g.). Three days later, mesenteric lymph nodes were excised, reduced to a single cell suspension by mechanical force, and stained for flow cytometry. CD45+, CD3+ CD4+ Vα2+, Vβ5+ cells were quantified for their proliferation cycles using FlowJo’s cell cycle tool and proliferation and division index were calculated.

### Statistics

Statistical analysis of all data was performed using Prism 10.0 (GraphPad, La Jolla, CA) with a Student’s t test or one-way or two-way ANOVA followed by post-hoc analysis with Tukey’s multiple comparison test. Individual data points are presented as mean ± standard error of the mean.

## Acknowledgements

These studies were funded NIH NIAID R01AI150647, R21AI148188 (CR), Animal Models of Infectious Disease Training Program T32AI060555 (MC), and a postdoctoral fellowship from Agencia Nacional de Ciencia y Desarrollo de Chile (ANID) #74200036 (V.T.R.). This project was supported by the UC Davis Flow Cytometry Shared Resource Laboratory with funding (NCI P30 CA093373) and technical assistance from Bridget McLaughlin, Jonathan Van Dye and Ashley Karajeh.

## Abbreviations

SP: Substance P
dpi: days post-infection
BEC: blood endothelial cells
LEC: lymphatic endothelial cells
ILC: Innate lymphoid cells
MLN: mesenteric lymph node
LN: lymph node
IFNγ: Interferon γ
IL-17: interleukin-17
IL-22: interleukin-22
FoxP3: forkhead box P3
iNOS/ Nos2: inducible nitric oxide synthase
LTi: lymphoid tissue inducer
NCR: natural cytotoxicity receptor
MAdCAM-1: mucosal addressin cell adhesion molecule
ICAM-1: intercellular cell adhesion molecule
VCAM-1: Vascular cell adhesion molecule
KO: knockout
cKO: conditional knockout
IEC: intestinal epithelial cells
TRPV1: transient receptor potential vanilloid 1
DC: dendritic cell
cDC: conventional dendritic cell
CFU: colony forming units
p.i.: post-infection
h.p.f: high power field
*C. Rod, C. Rodentium*: Citrobacter rodentium
LP: lamina propria
CP: CP96345
CGRP: Calcitonin gene related peptide
MHCII: major histocompatibility complex II.

---

**Figure S1.**
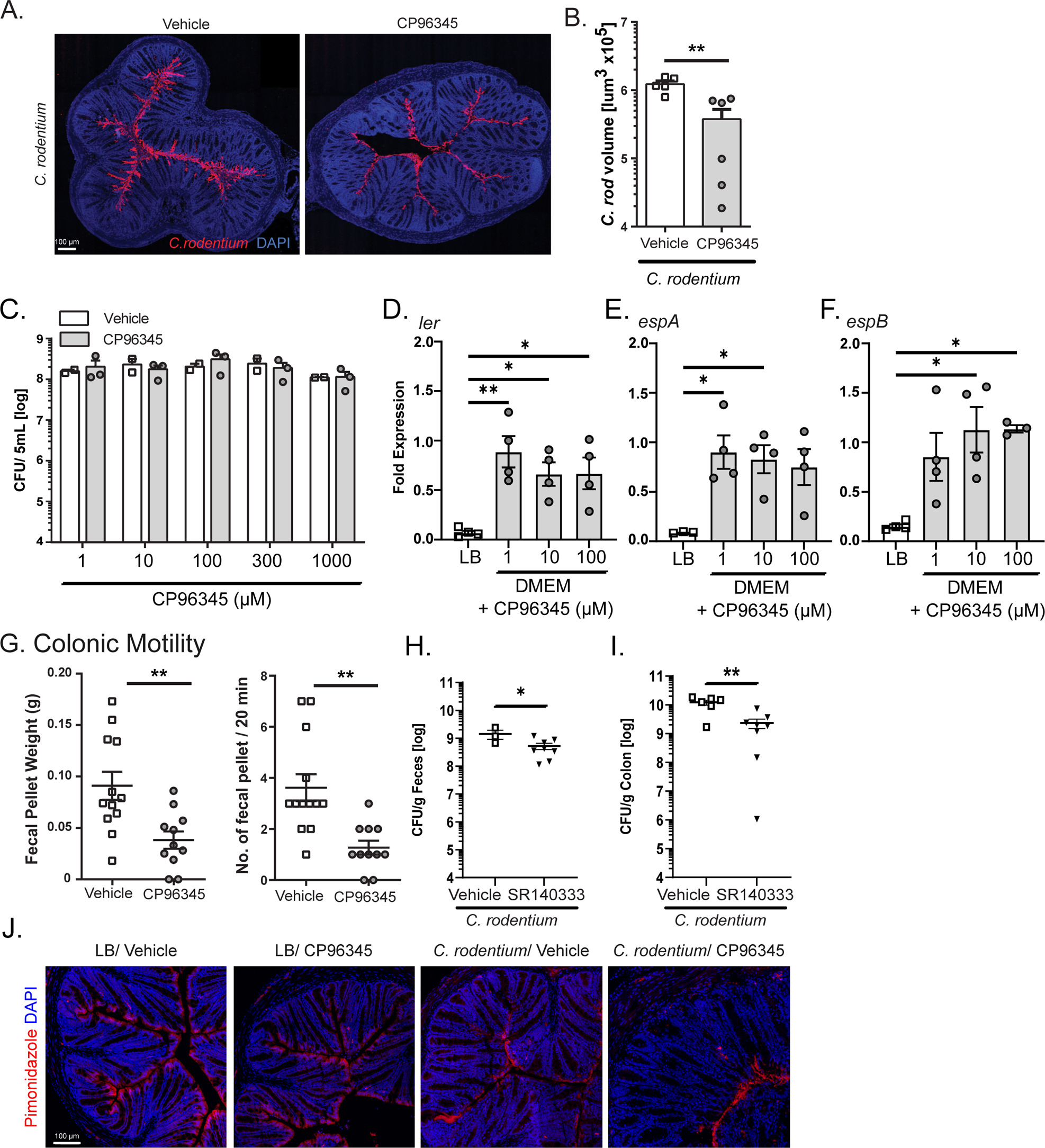
CP96345 reduces *C. rodentium* burden and GI motility without effecting bacterial viability or IEC oxygen availability. Immunofluorescence staining of colonic tissue sections with anti-*Citrobacter rodentium* specific antibodies were performed and images acquired with confocal microscopy **(A)** and the volume of stained bacteria quantified **(B)**. *C. rodentium* viability was quantified in the presence of CP96345 or vehicle (DMSO) in liquid culture by plating serial dilutions and enumerating CFU **(C)**. *C. rodentium* was cultured in LB or DMEM with increasing concentrations of CP96345 to assess expression of *ler* **(D)***, espA* **(E)**, and *espB* **(F)** mRNA was normalized to its equivalent percentage of DMSO control. Colonic motility was measured in mice treated with vehicle or CP96345 by fecal pellet weight and number of fecal pellets excreted in 20 minutes **(G)**. Fecal pellets **(H)** and colonic tissue **(I)** were quantified for *C. rodentium* burden 10 dpi in mice treated with vehicle or another TACR1 antagonist SR140333 intraperitoneally every two days. The degree of infection-induced oxygenation in the colonic tissues was determined by injection of Hypoxyprobe into uninfected and infected, vehicle or CP96345-treated mice 10 dpi. This probe was detected with immunofluorescence to detect the Hypoxyprobe adduct (pimonidazole), and images acquired by confocal microscopy **(J)**. Results are from individual mice or individual bacterial colonies, mean ± SEM, * = P ≤0.05, ** = P ≤0.01, *** = P ≤0.001. One-way ANOVA with Tukey’s post-hoc test was used. Luria broth (LB), *C. rodentium* (*C. rod*).

**Figure S2.**
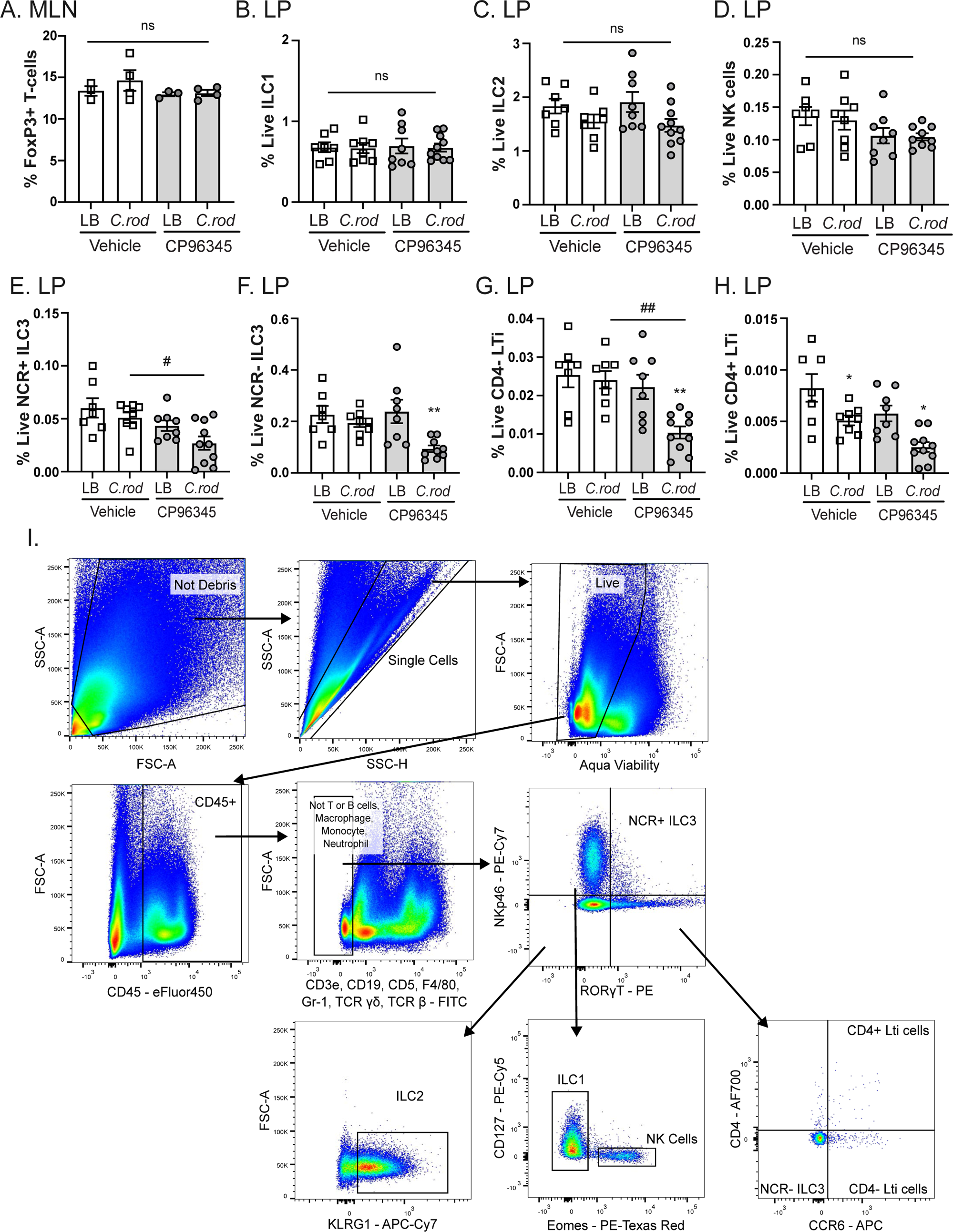
Certain subsets of ILCs are minimally impacted if at all with treatment of TACR1 antagonist. Mice were infected with *C. rodentium* or LB and treated with vehicle or CP96345 orally. Ten days post infection, MLN CD3+ CD4+ T-cells were assessed for their expression of FoxP3 via intracellular cytokine staining **(A)**. Three days post infection, colonic ILC1 **(B)**, ILC2 **(C)**, NK cells **(D)**, NCR+ ILC3 **(E)**, NCR-ILC3 **(F)**, CD4-Lti **(G)**, and CD4+ Lti **(H)** cells were quantified by frequency of live. Gating strategy for each of these populations **(I)**. Results are from individual mice, mean ± SEM, * = P ≤0.05, ** = P ≤0.01, *** = P ≤0.001 compared to uninfected controls of the same treatment and # = P ≤0.05, ## = P ≤0.01, ### = P ≤0.001 compared between treatment groups. One-way ANOVA with Tukey’s post-hoc test was used. Lamina propria (LP), Luria broth (LB), *C. rodentium* (*C. rod*), innate lymphoid cell (ILC), natural killer (NK), natural cytotoxicity receptor (NCR), lymphoid tissue inducer (LTi).

**Figure S3.**
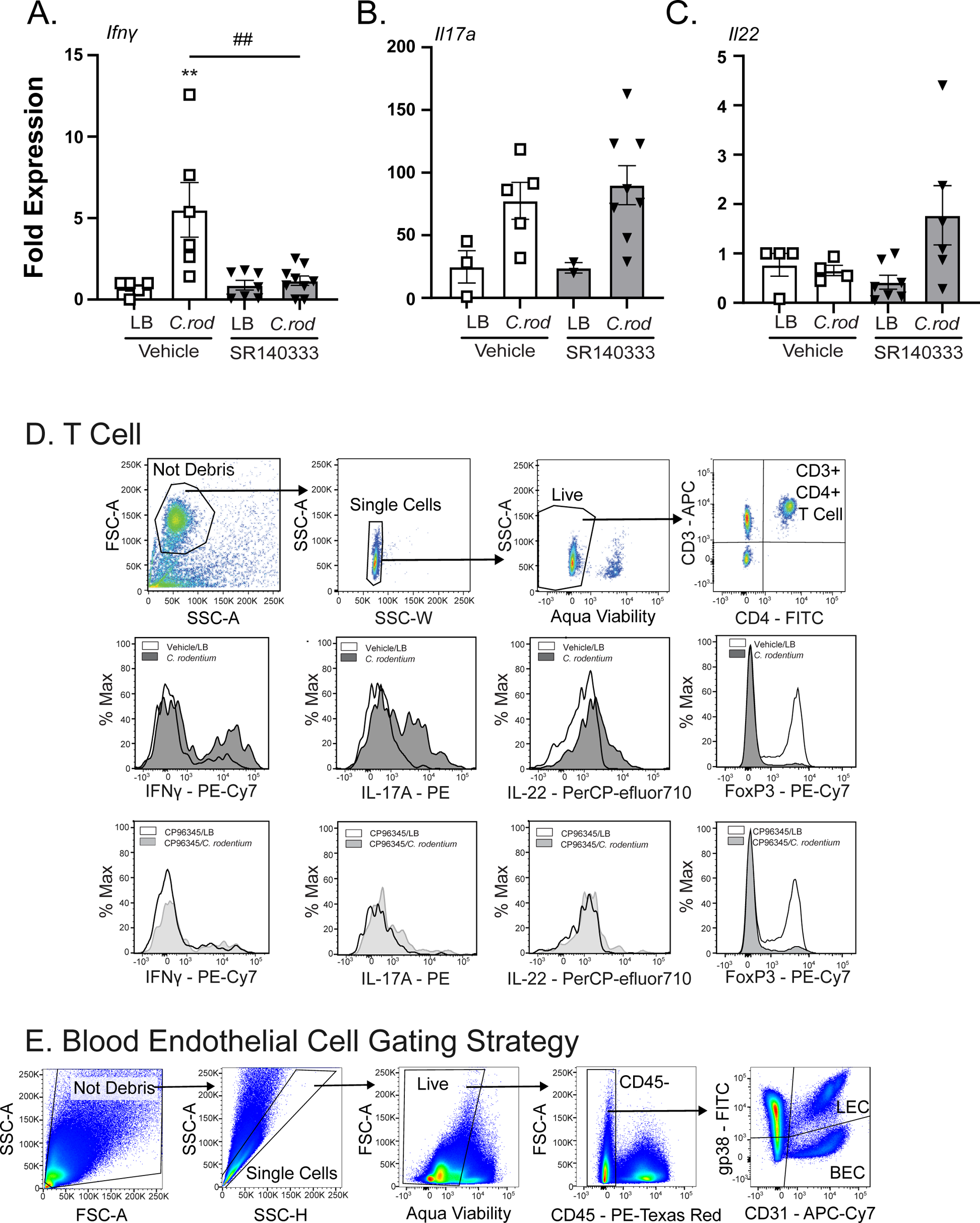
IFNγ expression is decreased in mice treated with different TACR1 antagonist through a separate drug delivery route & lamina propria flow cytometry gating strategies. Mice infected with *C. rodentium* or LB 10 dpi and treated with 1 mg/kg SR140333 i.p. or vehicle were assessed for their mRNA expression of *Ifnγ* **(A)**, *Il17a* **(B)**, and *Il22* **(C)** in colonic tissue. T-cell **(D)** and BEC **(E)** gating strategy used in Figure 2. **(A-C)** Results are from individual mice, mean ± SEM, * = P ≤0.05, ** = P ≤0.01, *** = P ≤0.001 compared to uninfected controls of the same treatment and # = P ≤0.05, ## = P ≤0.01, ### = P ≤0.001 compared between treatment groups. One-way ANOVA with Tukey’s post-hoc test was used. Luria broth (LB), *C. rodentium* (*C. rod*).

**Figure S4.**
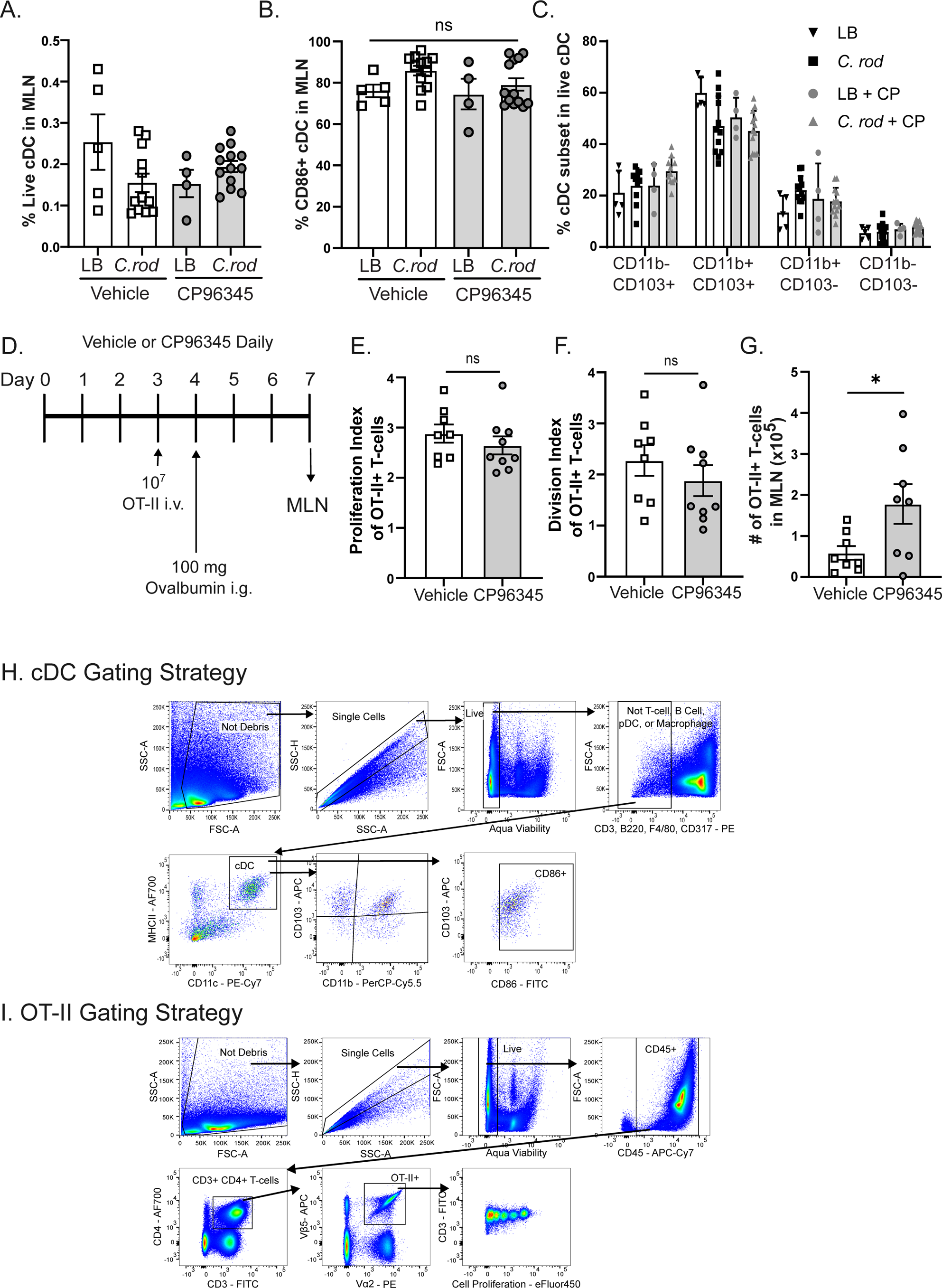
Conventional dendritic cell function and antigen specific T-cell proliferation is not reduced by TACR1 antagonism. Conventional dendritic cells (cDC) in MLN were analyzed from mice 10 dpi with *C. rodentium* or LB and treated with CP96345 or vehicle. The frequency of live cDC **(A)**, cDC expressing CD86 **(B)**, and different cDC subsets shown by their expression of CD11b and CD103 **(C)** were analyzed by flow cytometry. **(D)** WT mice were pre-treated for 3 days and continually treated for 4 days after with CP96345 or vehicle orally before having Cell Proliferation dyed CD4+ OT-II cells adoptively transferred. One day later, a bolus of ovalbumin was administered orally. Three days after that, MLN was analyzed for CD4+ OT-II+ T-cells. Proliferation index **(E)**, division index **(F)**, and total cell number of OT-II+ T-cells in the MLN **(G)** were quantified. Gating strategies for cDC **(H)** and OT-II T-cells **(I)**. Results are from individual mice, mean ± SEM, * = P ≤0.05, ** = P ≤0.01, *** = P ≤0.001. One-way ANOVA with Tukey’s post-hoc test was used. Luria broth (LB), *C. rodentium* (*C. rod*), CP96345 (CP).

**Figure S5.**
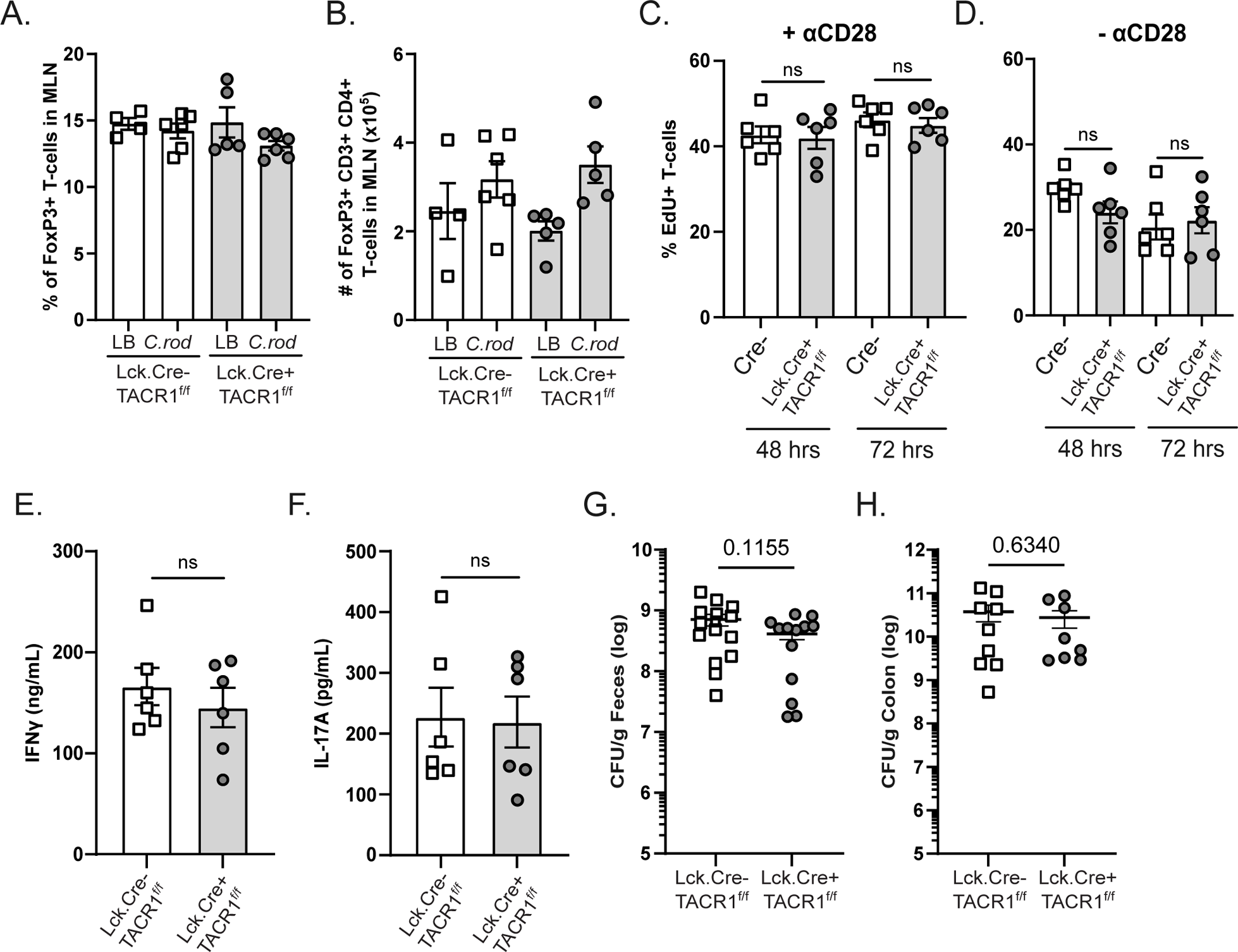
TACR1 deletion on T-cells does not cause significant defects of the T-cell’s ability to proliferate or produce cytokines *in vitro*. MLN were analyzed for FoxP3 expressing CD4+ T-cells in Lck.Cre+ TACR1^f/f^ mice and their Lck.Cre-TACR1^f/f^ littermates 10 dpi of *C. rodentium* infection or uninfected (LB) controls. Frequency of FoxP3+ in CD3+ CD4+ T-cells **(A)**, and total number of FoxP3+ CD3+ CD4+ T-cells in the MLN **(B)** were quantified. Splenic and lymph node CD4+ T-cells from Lck.Cre+ TACR1^f/f^ mice or Lck.Cre-TACR1^f/f^ littermates were cultured *in vitro* with anti-CD3 and anti-CD28 **(C)** or not **(D)** and assessed for their proliferation with EdU pulse for 2 hours or cultured in the presence of T-cell subset skewing cytokine cocktails to produce IFNγ **(E)**, or IL-17A **(F)**. Supernatant of these cultures were quantified by ELISA. Fecal **(G)** and colonic **(H)** *C. rodentium* bacterial burden was quantified in Lck.Cre+ TACR1^f/f^ mice and their Lck.Cre-TACR1^f/f^ littermates 10 dpi. Results are from individual mice **(A-B & G-H)** or from individual mice averaged between 2-5 technical replicates **(C-F)**, mean ± SEM, * = P ≤0.05, ** = P ≤0.01, *** = P ≤0.001. One-way ANOVA with Tukey’s post-hoc test was used. Luria broth (LB), *C. rodentium* (*C. rod*).

## Notes

### Competing Interest Statement

The authors have declared no competing interest.

